# Characterising neutrophil subtypes in cancer using human and murine single-cell RNA sequencing datasets

**DOI:** 10.1101/2023.07.13.548820

**Authors:** Rana Fetit, Mark White, Megan L. Mills, Xabier Cortes-Lavaud, Alistair McLaren, John Falconer, Kathryn Gilroy, Colin Nixon, Kristina Kirschner, Rene Jackstadt, Andrew D. Campbell, Owen J. Sansom, Colin W. Steele

## Abstract

Neutrophils are a highly heterogenous cellular population. However, a thorough examination of the different transcriptional neutrophil states, between health and malignancy, has not been performed. We utilised single-cell RNA-sequencing of human and murine datasets, both publicly available and independently generated, to identify neutrophil transcriptomic subtypes and their developmental lineages in health and malignancy. Datasets of lung, breast and colorectal cancer (CRC) were integrated to establish and validate the reproducibility of neutrophil gene-signatures. Pseudo-time analysis was used to identify genes driving neutrophil development from health to cancer. Finally, ligand-receptor interactions and signalling pathways between neutrophils and other immune cell populations in primary CRC and metastatic CRC were investigated. We define two main neutrophil subtypes in primary tumours: an activated subtype sharing the transcriptomic signatures of healthy neutrophils; and a tumour-specific subtype. This signature is conserved in murine and human cancer, across different tumour types. In CRC metastases, neutrophils are more heterogenous, exhibiting additional transcriptomic subtypes. Pseudo-time analysis implicates an IL1B/CXCL8/CXCR2 axis in the progression of neutrophils from health to cancer and metastasis, with effects on T-cell effector function. Assessment of global communication signalling identified CD4+ T-cells and macrophages as dominant regulators of the immunosuppressive, metastatic niche, whereas CD8+ T-cells are receivers of signals from other immune cells. We propose that the emergence of metastatic-specific neutrophil subtypes is driven by an IL1/CXCL8/CXCR2 axis, with the evolution of different transcriptomic signals that impair T-cell function at the metastatic site. Thus, a better understanding of the neutrophil transcriptomic programming could optimise immunotherapeutic interventions into early and late interventions, targeting different neutrophil subtypes.

## INTRODUCTION

Neutrophils are short lived cells released from the bone marrow in response to infection and inflammation and represent the most abundant white blood cells circulating. Traditionally thought of as terminally differentiated cells, neutrophils have been shown to demonstrate remarkable plasticity in response to different tissue environments(1), particularly to the tumour microenvironment(2). Indeed, in murine models of human disease, we observed significant phenotypic differences in response to inhibition of neutrophil populations using genetic and pharmacological approaches. In both metastatic pancreatic and colorectal cancer (CRC) models, we observed targeting neutrophil infiltration to metastases resulted in reduction of metastatic burden but with limited impact on primary tumour growth(3–5). These observations are in-keeping with other studies that have shown the potential of targeting metastasis associated neutrophils therapeutically in murine models of cancer(6,7). Understanding the regulation of neutrophils within metastases will permit future therapeutic efforts, with the aim of promoting an anti-tumoural neutrophil phenotype. Oncogenic *KRAS* driver mutations in lung and colorectal cancers(8,9) are thought to play a key role in driving a neutrophil phenotype within tumours, with clear upregulation of neutrophil chemotactic protein production from metastatic lesions(10). Systemic neutrophilia and inflammation have repeatably been associated with poor outcomes in CRC as well as in other cancers (11). Whilst these observations suggest the key role of neutrophils for promoting metastasis, dense neutrophil infiltration in the primary tumour microenvironment also correlates with poor prognosis in CRC suggesting neutrophils have roles in cancer progression at both primary and metastatic sites (11). Overall clear clinical and pre-clinical evidence exists for pro tumourgenic neutrophil populations(12).

Several studies have utilised single-cell RNA sequencing (ScRNAseq) to delineate the immune cell populations infiltrating the tumour microenvironment in different cancers and their respective mouse models, with a focus on macrophages and T-cell populations(13–18). However, to date, no study has thoroughly investigated neutrophil populations to specifically identify recurrent transcriptional subtypes in health and cancer. This underrepresentation of neutrophil populations in ScRNAseq datasets is largely owing to their short half-life, which dictates the fast processing of freshly procured samples, and the challenges in isolating adequate quality and quantity of RNA for downstream analysis, making neutrophils harder to capture using the common single-cell platforms. However, combining datasets can enable analysis of this underrepresented cell type that has not been extensively evaluated before. Likewise, neutrophil phenotypic differences between normal and tumour associated neutrophils have never been described at a single cell level.

Therefore, we sought to assess the differences in neutrophil transcriptional phenotypes between healthy tissue, primary tumour (PT) tissue and liver metastatic (LM) tissue across different cancer types: lung; breast; and CRC. Neutrophils have been shown in these cancer types to influence outcomes, in both mice and humans(19). We hypothesised that neutrophils show plasticity and adaptation to their surroundings to support anti- or pro-tumorigenic processes, with the metastatic site co-opting neutrophils to promote pro-tumorigenic neutrophil function. We demonstrated using publicly available ScRNAseq datasets and data generated from CRC murine models that two main subsets of neutrophils can be identified in health and cancer. We identified the developmental trajectory of these cells and observed a heterogenous group within LM tissue consistent with tissue specific adaptation at the metastatic site. This study lends novel insights to neutrophil single cell transcriptomic phenotypes and infers how these cells may be manipulated for therapeutic benefit in the future.

## MATERIALS AND METHODS

### Processing publicly available datasets

Datasets were retrieved from the GEOdatabase and National Omics Encyclopaedia(Table1) and processed using Seurat(version 4.3.0) on R(versions 3.17 and 4.1.1). Datasets were integrated by RPCA using the IntegrateData function then scaled and normalised. Dimension reduction was performed using PCA followed by clustering using the FindNeighbours and FindClusters functions. Marker genes for individual clusters were determined using the FindAllMarkers function and neutrophils were isolated by authors using the cluster identities and markers assigned in the original publications. Datasets were integrated to establish and test neutrophil gene-signatures using the AddModuleScore function. Pseudo-time analysis was performed using Slingshot(version 2.8.0) to identify neutrophil lineages. Gene expression along the different trajectories was performed using TradeSeq(version 1.14.0). Gene Set Enrichment (GSE), Gene Ontology (GO) and KEGG analyses were performed using ClusterProfiler(version 4.8.1) and EnrichR(version 3.2). Ligand-receptor (L-R) interactions and signalling pathways between neutrophils and other immune cell populations in primary and metastatic sites were investigated using CellChat (version 1.6.1). Software processing pipelines are listed in Table2 and all relevant code can be accessed on Github (https://github.com/ranafetit/NeutrophilCharacterisation).

**Table 1:**
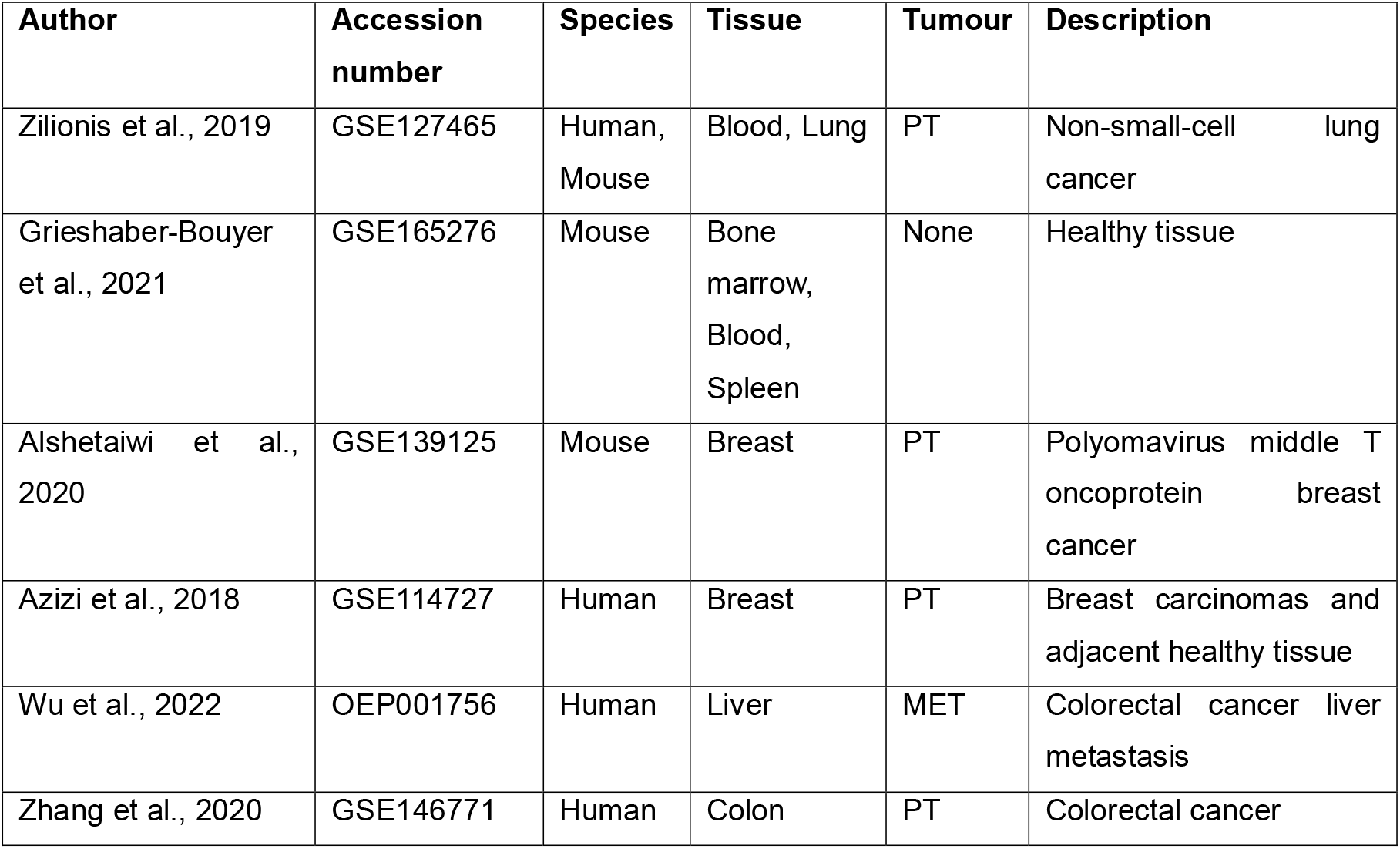
Description of public datasets used in this study.

**Table 2:**
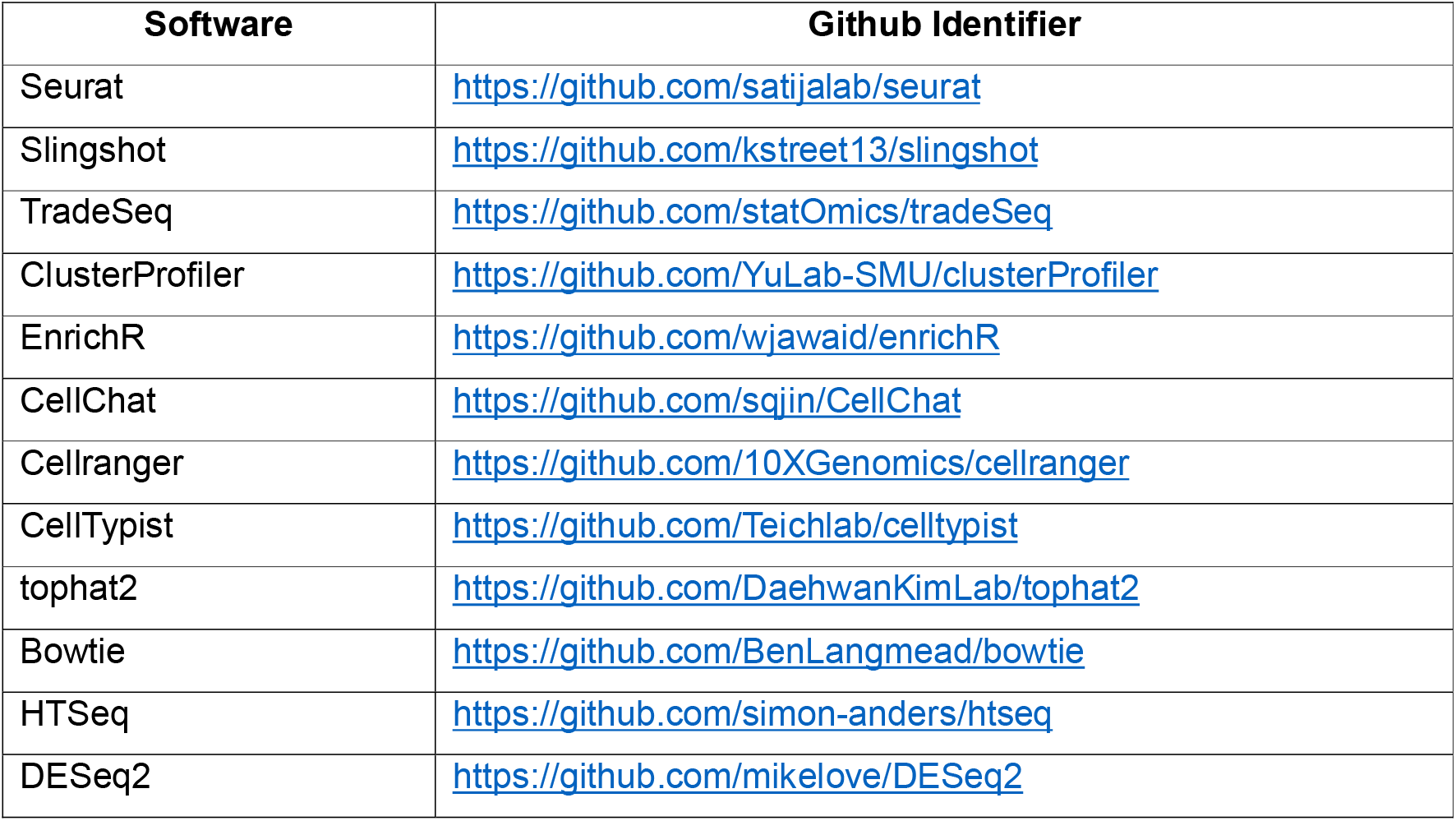
Links to software processing pipelines on Github.

| Software | Github Identifier |
| --- | --- |
| Seurat | https://github.com/satijalab/seurat |
| Slingshot | https://github.com/kstreet13/slingshot |
| TradeSeq | https://github.com/statOmics/tradeSeq |
| ClusterProfiler | https://github.com/YuLab-SMU/clusterProfiler |
| EnrichR | https://github.com/wjawaid/enrichR |
| CellChat | https://github.com/sqjin/CellChat |
| Cellranger | https://github.com/10XGenomics/cellranger |
| CellTypist | https://github.com/Teichlab/celltypist |
| tophat2 | https://github.com/DaehwanKimLab/tophat2 |
| Bowtie | https://github.com/BenLangmead/bowtie |
| HTSeq | https://github.com/simon-anders/htseq |
| DESeq2 | https://github.com/mikelove/DESeq2 |

### Mouse Housing and Ethics

All animal experiments were performed in accordance with the UK Home Office guidelines under project licences 70/9112 and PP390857, adhered to ARRIVE guidelines and were reviewed and approved by the University of Glasgow Animal Welfare and Ethical Review Board. Mice were housed in accordance with UK Home Office Regulations. Mice were fed standard chow diet and given drinking water ad libitum. A mixture of individually ventilated cages and conventional open top cages were used. Both genders of mice were used. Supplementary TableS1 summarises the numbers, sex and genotype of the mice used in this study.

### Mouse Models

The different intestinal cancer models are listed in Table 3. Two models of tumour genesis were used, aged genetically engineered mice and intracolonic transplants of murine derived organoids (Supplementary TableS1). All genetically engineered mouse models (GEMMs) were induced with a single 2mg intraperitoneal injection of Tamoxifen (Sigma-Aldrich T5648) when mice weighed >20g aged 6-18 weeks. Mice were aged until clinical endpoint defined as weight loss and/or hunching and/or piloerection and/or paling. For transplant mice, murine tumour derived organoids were injected intracolonically into male immune competent C57BL/6J mice (Charles River strain 632) using previously described methods(20). Tumour organoids were mechanically dissociated into fragments by pipetting and washed twice in PBS. Each mouse was injected with the equivalent of one well of a six well plate in 70uL of PBS. This was injected into the colonic submucosa using a Karl Storz TELE PACK VET X LED endoscopic video unit with associated needle. Transplanted mice were aged until clinical endpoint.

**Table 3:**
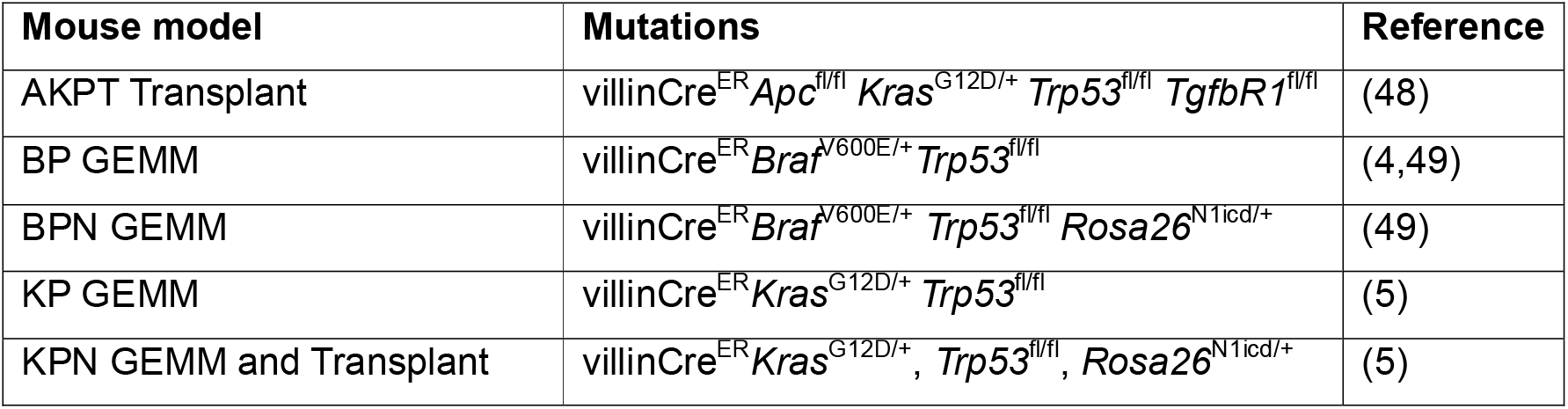
Description of CRC mouse models used.

### Tissue processing

Endpoint mice were culled and dissected. Entire primary tumour was removed and placed in PBS on ice. The tumour was then chopped into a smooth paste using a McIlwain Tissue Chopper. The paste was transferred to GentleMACS C tubes (Miltenyi Biotec, 130-093-237) with digestion enzymes from the Miltenyi Mouse Tumour Dissociation Kit (Miltenyi Biotec, 130-096-730) (2.35mL of RPMI1640, 100µL Enzyme D, 50µL Enzyme R, and 12.5µL Enzyme A). Samples were run on a GentleMACS Octo Dissociator with Heaters (Miltenyi Biotec, 130-096-427) using the 37C_m_TDK_1 programme. After digestion, samples were briefly spun, 10ml of RPMI-10%FBS-2mM EDTA was added and passed through a 70μm strainer. The resultant suspension was then spun down at 1800 RPM for 3 minutes at 4°C, supernatant discarded and the pellet resuspended in 0.5ml DPBS+0.05% BSA and transferred to a FACS collection tube on ice.

### ScRNA-sequencing

Mouse tumour cells were sorted using a BD FACSAria (BD Biosciences) and DAPI (Invitrogen, D1306) to remove dead cells, then loaded onto a Chromium Chip G using reagents from the 10x Chromium Single-Cell 3’ v3 Gel Bead Kit and Library (10x Genomics) according to the manufacturer’s protocol. Libraries were analysed using the Bioanalyzer High Sensitivity DNA Kit (Agilent Technologies) and sequenced on the Illumina NovaSeq 6000 with paired-end 150-base reads. Sequence alignment of single cell data to the mm10 genome was performed using the count tool from Cellranger(version6.1.2) according to the developers’ instructions, generating barcodes, features and matrix output files for each sample. Subsequent analysis was done using R (version 4.1.1) using Seurat (version4.0.4). Samples were input using the Read10X function, filtered to include cells with a minimum of 100 expressed genes and genes that are present in at least 3 cells, then further filtered to only include cells with <5% mitochondrial genes, <10% hemoglobin genes, >100 genes/cell and >400 reads/cell. Samples were then integrated by RPCA using the IntegrateData function before being scaled and normalised. Dimension reduction was performed using PCA followed by clustering using the FindNeighbours and FindClusters functions. Marker genes for individual clusters were determined using the FindAllMarkers function. Cell types were annotated using CellTypist and custom gene lists, and subset using the subset function.

### Bulk-RNA-sequencing of autochthonous CRC mouse models

Tissue processing, RNA isolation and sequencing were performed as described in(5). Briefly, primary and metastatic tumours were harvested from 5 villin-cre-ER, *Kras*^G12D/+^, *Trp53*^fl/fl^, *Rosa26*^N1CD/+^ (KPN) mice (model of metastatic CRC). Tumours from intestine, and liver were processed using the Mouse Tumour Dissociation Kit (Miltenyi Biotec #130-096-730) as per the manufacturer’s instructions, along with blood obtained by cardiac puncture upon terminal anaesthesia. Neutrophils were sorted based on CD48^-/lo^Ly6G^+^, CD11b^+^Ly6G^+^ expression and RNA was extracted using the RNeasy Mini kit (QIAGEN, #74104). Purified RNA quality was tested on an Agilent 2200 Tapestation using RNA screen tape. Libraries for cluster generation and RNA sequencing were prepared using the Illumina TruSeq RNA LT Kit after assessing RNA quality and quantity on Agilent 2200 Tapestation (D1000 screentape) and Qubit (Thermo Fisher Scientific), respectively. Libraries were run on Illumina Next Seq 500 using the High Output 75 cycles kit. Quality checks on the raw RNA-Seq data files were done using fastqc and fastq_screen (versions 0.11.2 and 0.11.3, respectively). RNA-seq paired-end reads were aligned to the GRCh38 mouse genome using tophat2 with Bowtie (versions 2.0.13 and 2.2.4.0, respectively). Expression levels were determined and analysed using HTSeq (version0.6.1) in R (version3.2.2), utilising Bioconductor data analysis suite and DESeq2.

### IHC of human CRCLM tissue

Access to colorectal cancer liver metastatic (CRCLM) patient tissue was authorized by the NHS Greater Glasgow and Clyde Biorepository under their NHS Research Ethics Committee (REC) approval with ethical approval granted in biorepository application #602. Upon successful metastatic liver resections, surplus tissue was stored in 4%PFA at 4^0^C for 20-48 hours. Samples were then transferred to 70% Ethanol and processed by standard histology processing techniques.

The following antibodies were stained on a Leica Bond Rx autostainer: CD3 (ab16669, Abcam) and TXNIP (40-3700, Thermo Scientific). All FFPE sections underwent on-board dewaxing (AR9222, Leica) and epitope retrieval using ER2 retrieval solution (AR9640, Leica) for 20 minutes at 95°C. Sections were rinsed with Leica wash buffer (AR9590, Leica) and peroxidase block was performed (Intense R kit; DS9263, Leica) for 5 minutes. Primary antibodies were added at optimal dilutions (CD3, 1/100; TXNIP, 1/400;) then rabbit envision secondary antibody (K4003, Agilent) was applied for 30 minutes. Sections were rinsed and visualised using DAB in Intense R kit.

FFPE sections for CD11b/ITGAM (49420, Cell Signaling) staining were loaded into the Agilent pre-treatment module for dewaxing and heat induced epitope retrieval (HIER) using high pH target retrieval solution (TRS) (K8004, Agilent). Sections were heated to 97□C for 20 minutes in high pH TRS buffer, rinsed in flex wash buffer (K8007, Agilent) then loaded onto the Dako autostainer. Peroxidase blocking (S2023, Agilent) was performed for 5 minutes. Primary CD11b/ITGAM antibody was added (1/400) for 35 minutes, then rabbit envision secondary antibody was applied for 30 minutes. Sections were rinsed before applying Liquid DAB (K3468, Agilent) for 10 minutes. Sections were washed in water and counterstained with haematoxylin z (RBA-4201-00A, CellPath). Finally, all sections were rinsed in tap water, dehydrated through graded ethanol’s, placed in xylene then coverslipped using DPX mountant (SEA-1300-00A, CellPath).

## RESULTS

### Neutrophils exhibit distinct tissue-specific and tumour-specific signatures

To examine the transcriptomic signatures of neutrophil subtypes in healthy and tumour tissue, we integrated neutrophil clusters from bone marrow, blood, lung and spleen of healthy mice(14), together with neutrophils from tumour-bearing mouse models of non-small cell lung cancer (NSCLC)(13) and CRC (KPN). Both NSCLC and CRC tumour models shared a comparable C57BL/6 background with *Kras* and *Trp53* mutations. For CRC, neutrophils were derived from two models of tumour genesis: aged GE mice and intracolonic transplants of murine derived organoids, with the majority being from the latter (Fig.S1A). Unsupervised clustering of neutrophil transcriptomic signatures revealed distinct neutrophil clusters based on their tissue of origin in health (Fig.1A). KPN and lung adenocarcinoma neutrophils formed distinct tumour-specific clusters, suggesting transcriptomic differences (Fig.1B).

**Fig.1.**
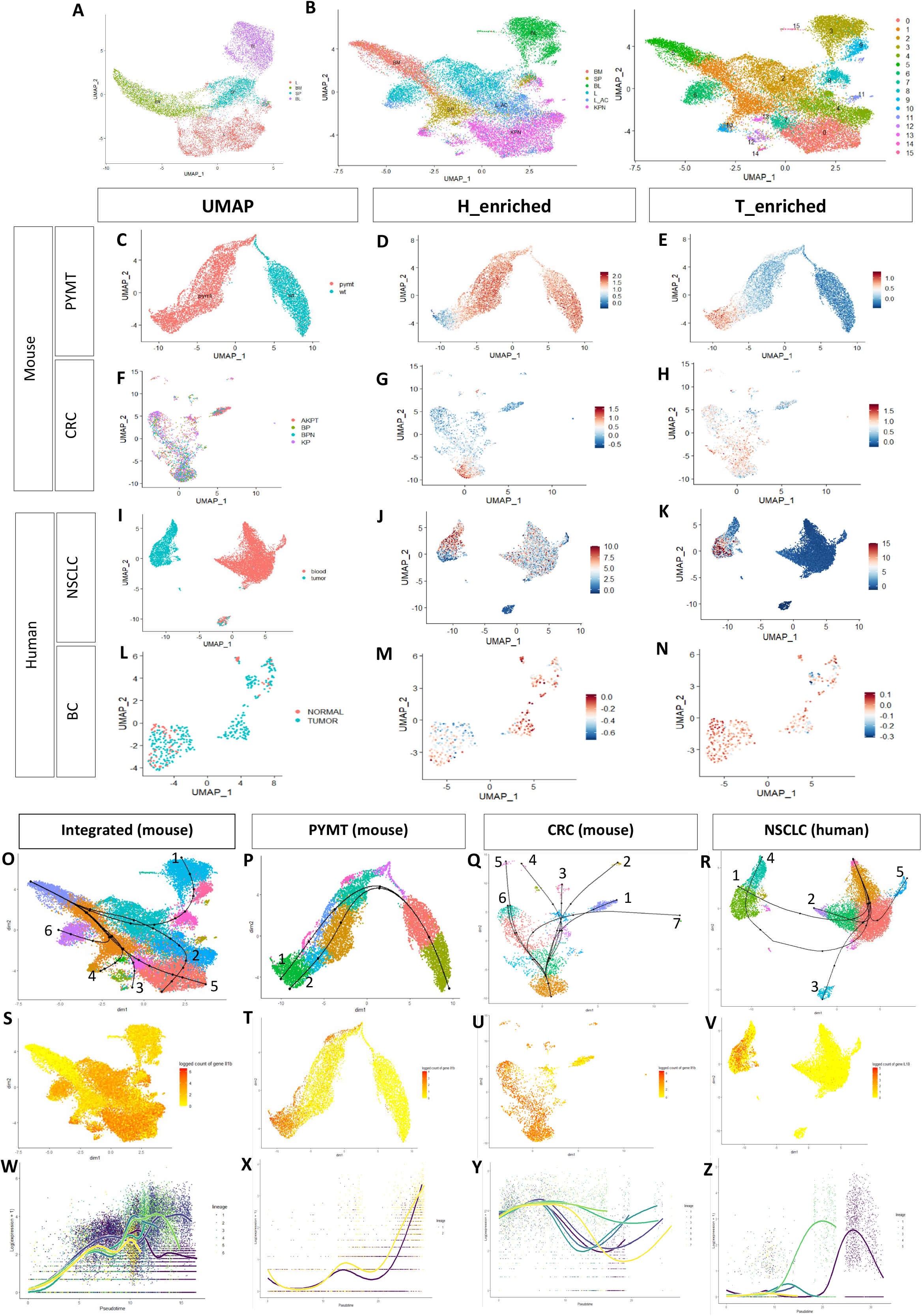
Characterisation of neutrophil signatures and lineages in health and primary tumours. (A) UMAP plot of healthy neutrophils grouped by tissue type. BM: healthy bone marrow, SP: healthy spleen, L: healthy lung, BL: healthy blood. (B) UMAP plots of healthy and tumour-derived neutrophils grouped by tissue type and Seurat clusters (0-15). L_AC: lung adenocarcinoma, KPN: colorectal cancer with *Kras*, *Trp53* and Notch mutations. (C) UMAP plot of neutrophils in mouse breast cancer model. WT: healthy breast tissue, PYMT: polyomavirus middle-T oncoprotein tumour. (D,E) Scoring of Healthy_enriched (H_enriched) and Tumour_enriched (T_enriched) neutrophil signatures. (F) UMAP plot of neutrophils in mouse CRC model. All neutrophils are tumour-derived. (G,H) H_enriched and T_enriched signatures in CRC. (I) UMAP plot of human NSCLC neutrophils. Blood: blood-derived, tumour: tumour-derived. (J,K) H_enriched and T_enriched signatures NSCLC. (L) UMAP plot of human breast cancer (BC) neutrophils. Most neutrophils are tumour derived. (M,N) H_enriched and T_enriched signatures in BC. (O-R) Unsupervised pseudo-time analysis of neutrophils in mouse and human datasets. Lineages in the individual datasets are numbered. (S-V) *Il1b/IL1B* is differentially expressed at the end of tumour-specific lineages. (W-Z) Estimated smoothers for *Il1b/IL1B* expression over pseudo-time across the different lineages.

KPN neutrophils encompassed clusters: 0,4,7,8,10,11,12 and 14 (Fig.1B and Fig.S1B, Fig.S2A). Clusters 0 and 7 were enriched for *Cxcl2* and *Thbs1*, which encode proteins that influence neutrophil motility and chemotaxis. Cluster 0 also expressed *Ccl4* and *Ccl3*, critical for T-cell recruitment and antitumor immunity(21) (Supplementary Table S1). Cluster 10 expressed *Cd74*, which plays a role in neutrophil accumulation(22). Cluster 4 was common to both KPN and NSCLC primary tumours (PTs) (Fig.1B; Fig.S1B), and expressed *Cdkn1a, Ppia, Gngt2 Ier3,* and *Rps27l* (Supplementary Table S2). Three smaller clusters were shared between both PTs: Cluster 12, enriched for the lysosomal genes *Lyz1* and *Psap*, Cluster 14 expressing *Ppia, Jun* and *Slfn4*, and Cluster 11 enriched for *S100a10* and *Ptma.* Finally, Cluster 8 was equally conserved across healthy and tumour-associated neutrophils (Fig.1A,B; Fig.S1B) and was enriched for interferon (IFN) markers: *Isg15, Rsad2, Ifit3, Ifit1* and *Slfn4* (Supplementary Table S2), a phenomenon previously reported in several ScRNAseq studies of neutrophils(23).

### Neutrophils in PTs encompass two transcriptional subtypes in mice

Using the highly expressed markers in healthy and tumour tissues (Supplementary Table S2), we defined two neutrophil signatures. The first represented neutrophils associated with healthy tissue (Healthy-enriched; (H_enriched). This subtype was observed in both healthy and tumour tissue. The second signature was specific to tumour-associated neutrophils (Tumour-enriched; T_enriched) (Fig.1C-N).

To validate the established signatures across different tissues and tumours, we scored them on additional neutrophil ScRNA-seq datasets derived from a mouse model of healthy and breast cancer tissue, utilising the mouse mammary tumour virus (MMTV) promoter–driven expression of the polyomavirus middle-T oncoprotein (PYMT, GSE139125) and neutrophils derived from a compendium of CRC mouse models generated in our lab (Table 2).

In PYMT, neutrophils clearly separate into distinct healthy and tumour-specific clusters, recapitulating the findings in NSCLC and CRC datasets (Fig.1C,F). Signature scoring in both PYMT and CRC mouse models revealed that within the PT, tumour-specific neutrophils can be separated into two subgroups: (1) activated neutrophils, which are transcriptionally similar to neutrophils from healthy tissue (Fig.1D,G), and (2) a subtype specific to PTs (Fig.1E,H). Both signatures were preserved in both GEM and transplant models of CRC (Fig. S2B,C).

### Neutrophil signatures are conserved between mouse and human

We then investigated whether these signatures (Table 4) could be translated to humans, using 2 datasets: patient-derived neutrophils from NSCLC tumour and blood (Fig. 1I, <u>GSE127465</u>); and breast carcinomas (BC) and adjacent healthy tissue (Fig.1L, <u>GSE114727</u>). Signature scoring in NSCLC confirmed the presence of both neutrophil subsets within the PT (Fig.1J,K) recapitulating the trends observed in mice. Blood-derived neutrophils largely resemble the H_enriched subtype (Fig. 1J). Although the BC dataset contained very few cells, we successfully observed the enrichment of both neutrophil subtypes in tumour-derived neutrophils (Fig.1M,N). Our analysis validates the presence of both neutrophil transcriptomic subtypes in patient PTs, albeit to different extents in the different cancer types and tissues, implying a role of the neutrophil’s environment in shaping their transcriptome.

**Table 4:** Healthy- and Tumour enriched neutrophil signatures in human and mouse.

| Human |  |  | Mouse |  |  |
| --- | --- | --- | --- | --- | --- |
| Healthy-enriched | Tumour-enriched | Metastasis-enriched | Healthy-enriched | Tumour-enriched | Metastasis-enriched |
| MMP8 | CDKN1A | TXNIP | Mmp8 | Cdkn1a | Txnip |
| IFITM2 | PPIA | RIPOR2 | Ifitm6 | Ppia | Riopr2 |
| IFITM3 | IFITM1 | CXCR2 | S100a6 | Ifitm1 | Cxcr2 |
| S100A6 | TAGLN2 | FKBP5 | Lyz1 | Tagln2 | Fkbp5 |
| LYZ | ISG15 | CEBPD | Lyz2 | Isg15 | Cebpd |
| CTLA4 | GNGT2 | STK17B | Ctla2a | Gngt2 | Stk17b |
| CHI3L1 | CXCL8 | SMAP2 | Chil3 | Cxcl12 | Smap2 |
| G0S2 | CCL4 | CTSS | G0s2 | Ccl4 | Ctss |
| FPR2 | CD14 | JAML | Fpr2 | Cd14 | Jaml |
|  | RPS27L | IGHM |  | Rps27l | Ighm |
|  | IER3 | CD74 |  | Ier3 | Cd74 |
|  | CCL3 |  |  | Ccl3 |  |
|  | IFIT3 |  |  | Ifit3 |  |
|  | IFIT1 |  |  | Ifit1 |  |
|  | IL1B |  |  | Il1b |  |
|  | WFDC1 |  |  | Wfdc17 |  |
|  | THBS1 |  |  | Thbs1 |  |
|  | PTMA |  |  | Ptma |  |

### Pseudo-time analysis demonstrates neutrophil lineages progress from H_enriched towards T_enriched neutrophils

To investigate the developmental trajectory of neutrophils from health to cancer, we performed unsupervised pseudo-time analysis on our integrated mouse dataset (Fig.1O). Individual lineages are shown in Supplementary Table S3. Our results recapitulated the Neutrotime lineage(14): from bone marrow to spleen and blood in healthy neutrophil populations (Fig.1O; Lineage1, Supplementary Table S3), with additional lineages as tumour-specific clusters develop. This trend was equally observed in the murine PYMT and CRC datasets (Fig.1P,Q). In human NSCLC, lineages begin from the blood-derived clusters enriched for the H_enriched signature, and progress towards the tumour-derived clusters enriched for the T_enriched signature (Lineages 1 and 4, Supplementary Table S3 and Fig.1R). Our data identified a developmental trajectory beginning with activated, healthy neutrophils and ending at tumour-specific neutrophils in these datasets.

### Interleukin-1B (*Il1b*) is a driver of T_enriched Neutrophil signature

We analysed gene expression along the different trajectories to identify genes that drive neutrophil differentiation from health to cancer-associated lineages. Specifically, we investigated the genes differentially expressed at the end of the lineages compared with the start. In our integrated dataset, *Il1b* was upregulated in the lineages ending with the tumour clusters (Fig.1S,W). *Il1b* was also among the top 30 lineage-specific differentially expressed genes and was specific to the T_enriched neutrophil clusters PYMT (Fig.1T,X; Supplementary Table S4). This was true for CRC (Fig. 1U,Y; Lineages 4 and 6; Supplementary Table S5). In human-NSCLC, the same trend was observed in Lineages 1, 2 and 4 (Supplementary Table S6; Fig.1V,Z). The human BC dataset was too small to perform such an analysis. Taken together, our lineage-specific differential gene expression analyses implicate *Il1b* in the progression of neutrophils towards the T-enriched population in PT.

### Neutrophils in CRC liver metastasis (LM) display heterogenous transcriptional programmes

To investigate whether these neutrophil subtypes are present in metastatic cancer, we isolated neutrophils from the publicly available CRCLM dataset(16) (Fig.2A) and scored them for the two signatures we established in PTs. Neutrophils in CRCLM expressed both H_and T_enriched signatures. However, a remarkable overlap between the two signatures was observed, with no clear separation between the two subtypes, reflecting their heterogeneity (Fig.2B). Some clusters were not enriched for either the H_ or T_enriched neutrophil signature (Fig.2B, blue arrow), suggesting the presence of an additional transcriptionally segregated neutrophil population, specific to metastatic CRC.

**Fig.2.**
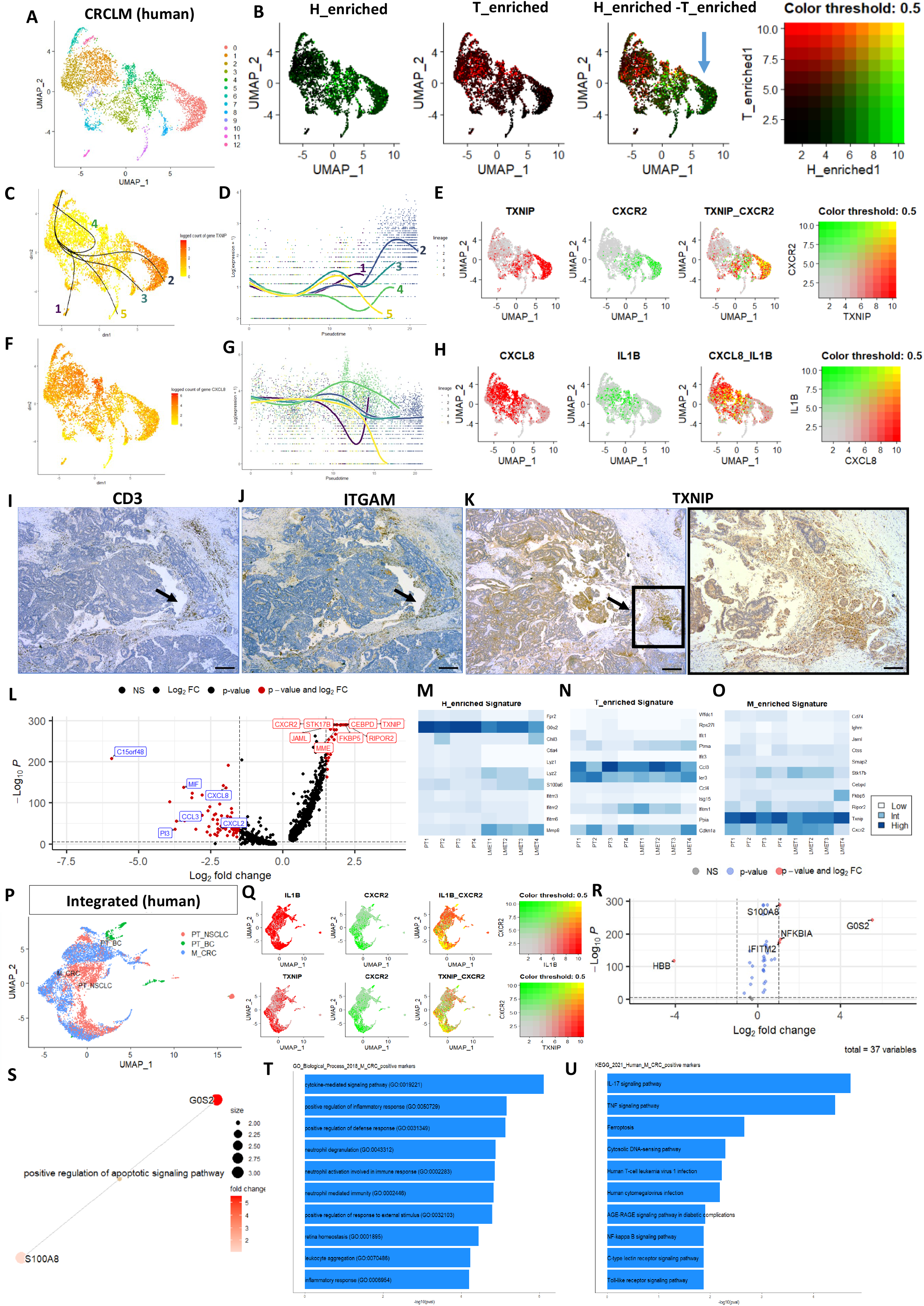
Characterisation of neutrophils in metastasis. (A)UMAP of neutrophils in CRCLM. (B) Co-expression of H_enriched and T_enriched signatures. One cluster is not enriched for either signature (blue arrow). (C,D) Unsupervised pseudo-time analysis and estimated smoothers for TXNIP expression over the different numbered pseudo-time lineages. (E) Co-expression of TXNIP and CXCR2. (F,G) Expression and estimated smoothers for CXCL8 over pseudo-time. (H) Co-expression of CXCL8 and IL1B. (I,J) IHC staining of CD3 (T-cells) and ITGAM (Neutrophils) in a patient CRCLM sample at 4x, scalebars=50µm. Black arrows indicate regions where immune cells cluster. (K) IHC staining of TXNIP in a patient CRCLM sample at 4x (left) and 10x(right). Scalebars=50µm. (L) Differentially expressed genes in metastasis-specific neutrophil cluster. (M-O) H_enriched, T_enriched and M_enriched gene signatures in mouse Bulk-RNAseq neutrophil dataset. PT: Primary tumour, LMET: Liver metastasis. (P)UMAP plot of integrated human neutrophils from primary tumours (PT) and metastatic (M) datasets of different cancers. (Q) Co-expression of *CXCR2* with *IL1B* (top) and *TXNIP* (bottom). (R) Differential gene expression between neutrophils in malignancy compared to PT. (S-U) GESA, GO and KEGG analysis of M_CRC neutrophils.

Unsupervised pseudo-time analysis revealed 5 neutrophil lineages, starting from the clusters enriched for the T_enriched signature (Fig.2C). All lineages shared the same sequence for the first 6 clusters and differed at their terminal clusters (Supplementary Table S3). We focused on lineages 2 and 4 because lineage 2 progressed towards the cluster not enriched for either signature observed in PT while lineage 4 terminated with a cluster expressing both signatures observed in PT (Supplementary Table S3). Collectively, our findings suggest progressive transcriptomic development of neutrophil phenotype from healthy to tumour specific signatures in PT and finally, a metastatic-specific neutrophil subtype.

### CRCLM-specific neutrophils display T-cell suppressive markers

To characterise the transcriptomic signature of the metastatic-specific neutrophil population, we identified marker genes for the lineage endpoints. The endpoint of Lineage 4 (Cluster 5; Supplementary Table S3) was enriched for the chemokine *CXCL8* (Fig.2F,G), the major ligand for G-Protein coupled receptor CXCR2 and associated with immune suppression and tumour progression in this context(24). This cluster also highly expressed *IL1B* (Fig.2H), supporting our hypothesis that *IL1B* is implicated in the progression of neutrophils towards malignancy associated phenotypes. Lineage2 endpoint was enriched for the mRNA encoding Trx-interacting protein (*TXNIP*, Fig.2C,D), the upregulation of which inhibits TRX1 and restrains late T-cell expansion(25). This cluster was also enriched for the chemokine receptor CXCR2 (Fig.2E), which is a commonly studied target in murine models of cancer influencing metastatic burden, suggesting these neutrophils identified are functionally relevant. We confirmed the expression of TXNIP in a patient CRCLM sample (Fig. 2K), in tumour regions where immune cells cluster (Fig. 2I-J; black arrows). Moreover, among the 10 most highly expressed markers in this cluster were the genes: *RIPOR2* and *STK17B*, which are important for naïve T-cell quiescence, survival and activation(26,27) (Fig.2I; Supplementary Table S7). Our findings suggest that the metastasis-specific neutrophil subtype transcribes genes that are T-cell suppressive.

### Murine neutrophils express a metastasis-specific signature in CRCLM bulk-RNA-seq dataset

We selected the top 11 highly expressed genes in the metastasis-specific neutrophil clusters, which we called Metastasis-enriched (M_enriched) signature (Table 3; Fig.2L,O; Supplementary Table S7). We then compared the established H_enriched, T_enriched and M_enriched gene signatures in a bulk-RNA-seq dataset generated from neutrophils from an autochthonous KPN mouse model of CRC-PT and LM generated in our lab. Higher expression of H_enriched genes was observed in LM compared with the PT tissue (Fig.2M). Neutrophils in both PT and LM tissue equally expressed the T_enriched signature, with a higher expression of few genes such as *Ptma* and *Ifitm1* in LM (Fig.2N). A similar trend was observed in the M_enriched signature, where an increase in *Stk17b, Fkbp5, and Cxcr2* was observed in LM (Fig.2O). This further validates the trends observed in our ScRNAseq analysis and demonstrates cross-species relevance.

### GSE analysis implicates IL-17/CXCR2 axis in metastatic neutrophil populations

We integrated patient-derived neutrophil Sc-RNAseq signatures from lung and breast cancer PTs (PT_NSCLC and PT_BC), with neutrophils from CRCLM tissue (M_CRC) (Fig.2P). Co-expression analysis revealed that neutrophils from both PT and LM tissue expressed *IL1B*, however, *CXCR2* expression was largely specific to LM (Fig.2Q). Metastatic neutrophils co-expressed CXCR2 and TXNIP (Fig.2Q), highlighting the specificity of these markers to neutrophils in CRCLM. Differential gene expression between neutrophils in PT and in CRCLM revealed the upregulation of the NETosis marker *G0S2* and *NFKB1A*. (Fig.2R). Supplementary Table S8 shows the top10 differentially expressed genes, grouped by tumour type. Using the differentially expressed genes in M_CRC neutrophils (Supplementary Table S9), we performed GSE analysis. *G0S2* and *S100A8* were upregulated and are implicated in positive regulation of apoptotic signalling (Fig.2S). GO analysis revealed an upregulation of cytokine-mediated signalling and positive regulation of inflammatory response (Fig.2T). KEGG analysis revealed the up regulation of IL17, TNF and NF-kappa-beta signalling pathways (Fig.2U). This supports the data implicating the IL17/CXCR2 axis in metastatic neutrophil populations(28).

### CRCLM-derived CD4+T-cells transcriptionally diverge from their PT counterparts

We revisited the publicly available datasets of CRC PT(17) and CRCLM(16) to isolate T-cells and investigate transcriptomic differences in metastasis given the T-cell suppressive phenotype of neutrophils found in CRCLM. Upon integration, CD8+T-cells from both PT and LM largely co-cluster, reflecting their transcriptomic similarity. However, CRCLM-derived CD4+T-cells formed a distinct cluster (Fig. 3A,B). Differential gene expression analysis revealed 660 and 26 differentially expressed genes for the CRCLM-derived CD4 and CD8+T-cells, respectively, compared with their equivalent PT populations (Supplementary Tables S10 and S11). Henceforth, we focused on CD4+T-cells. *IL1B* and *CXCL8* were upregulated, the same genes we identified as drivers of metastatic neutrophil subtype (Fig.3C), suggesting that the IL1B/CXCL8/CXCR2 axis drives the interaction between neutrophils and T-cells in metastatic tissue.

**Fig.3.**
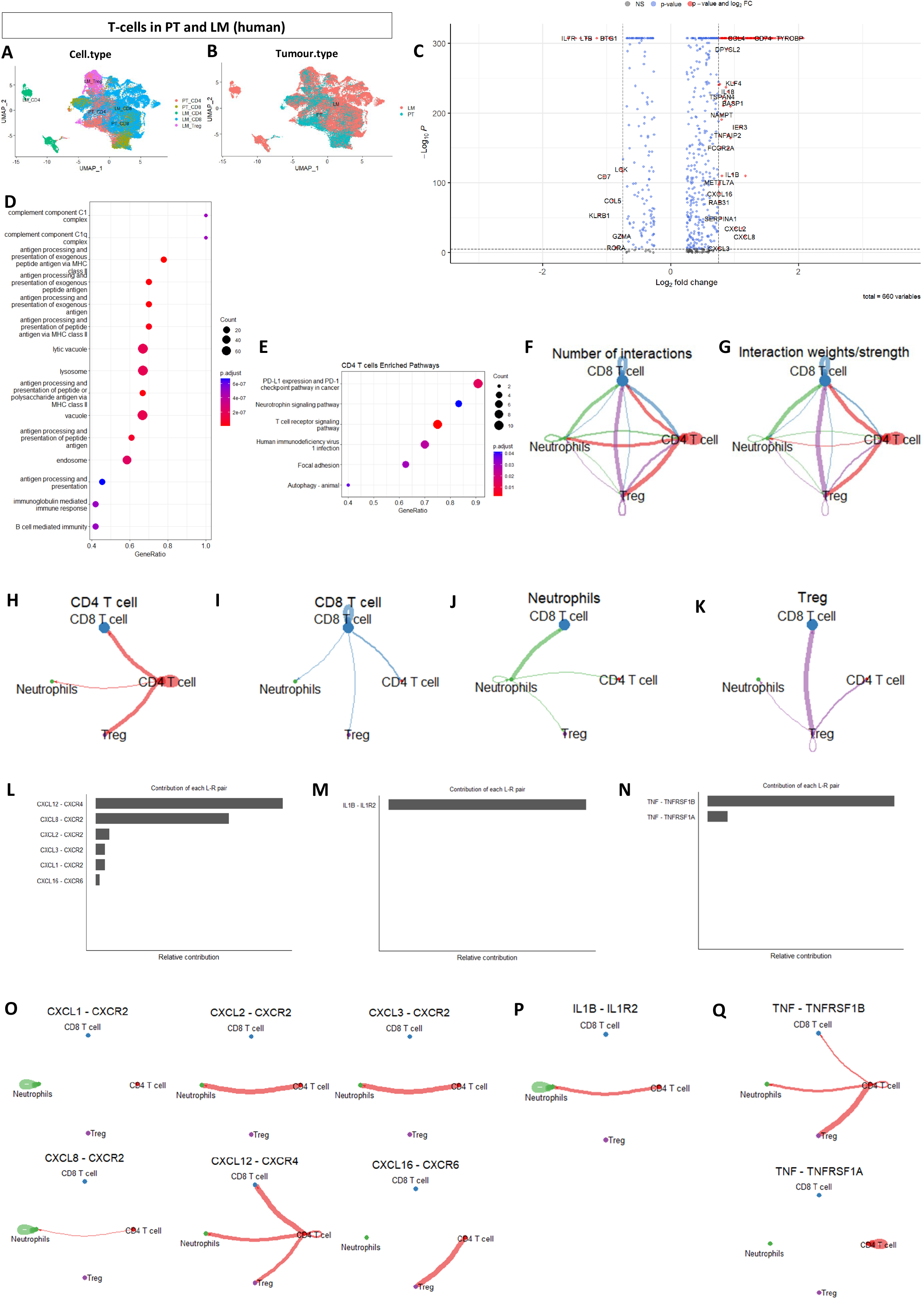
CD4+T-cells are transcriptomically altered in metastasis. (A,B)UMAP plots of CD4+ and CD8+T-cells in PT and LM grouped by cell type and tumour type, respectively. (C) Differential gene expression of CD4+T-cells in LM compared to PT. (D,E) GO and KEGG analyses of metastatic CD4+T-cells. (F,G) Global cell-cell communication network and the interaction strengths between Neutrophils, CD4+,CD8+ and Treg cells. (H-K) Outgoing signals sent from each cell group. (L-N) The contribution of each L-R pair to the overall signalling pathway for CXCL, IL1 and TNF pathways. (O-Q) Visualization of the cell-cell communication patterns mediated by each significant L-R pair for the three pathways.

In addition, we observed a downregulation of *RORA*, implicated in CD4 T-cell activation(29), together with Granzyme A (*GZM*A) and pro-inflammatory lipid-mediator leukotriene B (*LTB*); further suggesting the possibility of impaired cytotoxic CD4+T-cell function in CRCLM. GO and KEGG analyses revealed the dysregulation of biological processes converging on the complement system, antigen processing and presentation, together with perturbations in PD1-PDL1 and T-cell receptor signalling pathways in the CRCLM-derived CD4+T-cells (Fig.3D,E). Collectively, this suggests that the regulatory function of CD4+T-cells may be impaired in metastasis, contributing to an immunosuppressive phenotype.

### Neutrophils and CD4+T-cells interact through IL1, CXCL and TNF signalling pathways in CRCLM

We then assessed the global cell-cell communication network to investigate how the impaired CD4+T-cells in the metastatic niche may influence neutrophils and other T-cell subtypes. Signals from CD4+T-cells interact with CD8+T-cells and T-regulatory cells (Tregs) and, to a lesser extent, neutrophils (Fig.3F-H). Outgoing signals from neutrophils are received by CD8+T-cells (Fig.J), supporting our hypothesis that neutrophils impair CD8+T-cell function in metastasis. Signals from CD8+T-cells are mostly autocrine (Fig.3I) and Tregs mainly influence CD8+T-cells (Fig.3K). We identified 20 signalling pathways showing significant communications between neutrophils, CD4+, CD8+ T-cells and Tregs (Supplementary Fig.S3). Supplementary Fig.S4A-D show the significant ligand-receptor (L-R) interactions between neutrophils, CD4+T-cells, CD8+T-cells and Tregs to other target cell groups. Neutrophils primarily communicate with CD8+T-cells through the MHC-I pathway (Fig.S4A). CD4+T-cells strongly interact with CD8+T-cells through MHC-I and MHC-II pathways and with Tregs through the MHC-II pathway (Fig.S4B). CD8+T-cells communicate with CD4+T-cells and neutrophils through CD45 and ANNEXIN signalling pathways, respectively (Fig.S4C). Finally, Tregs target CD8+T-cells through the MHC-I signalling pathway (Fig.S4D).

We then focused on IL1, CXCL and TNF pathways, based on their involvement in defining the phenotype of metastatic neutrophils. CXCL12-CXCR4 and CXCL8-CXCR2 were the major L-R interactions observed(Fig.3L). CD4+T-cells are major senders of the CXCL12-CXCR4 signals, whereas CXCL8-CXCR2 signals are largely autocrine within neutrophils (Fig.3O). IL1B-IL1R2 significantly contributed to the IL1 pathway in CRCLM (Fig.3M), an L-R interaction largely driven by the CD4+T-cells and neutrophil interactions, as well as neutrophils’ autocrine signalling (Fig.3P). This observation supports our pseudo-time findings in the different neutrophil populations. Finally, CD4+T-cells communicate with Tregs, neutrophils and CD8+T-cells through TNF-TNFRSF1B interactions (Fig.3N,Q). Our findings suggest that within the metastatic niche, neutrophils primarily target CD8+T-cells through the MHC-I pathway, in addition to their autoregulation through CXCL and IL1 pathways with CD4+T-cells undertaking a prominent regulatory role.

### CD4+T-cells are dominant signal senders in CRCLM

To elucidate how cells coordinate different pathways to drive communication, we investigated the global communication patterns between the different immune cell populations. We identified 4 outgoing patterns (Fig.4A) and 4 incoming patterns (Fig.4B). Outgoing signals from CD4+T-cells formed the largest communication pattern, with MHC-II and PECAM signalling being autocrine (Fig.4A,B). Outgoing signals from CD8+T-cells converge on the CLEC pathway, which is autocrine, together with signals from ANNEXIN, CD99 and IFN-II pathways. Tregs send signals along the LCK and VCAM pathways and are recipient to CD86 signalling. The IL1 signalling pathway is the most prominent outgoing pathway for neutrophils, which is also autocrine (Fig.4A,B). Neutrophils are influenced by CXCL ligands and ICAM signals from CD4+T-cells, ANNEXIN signals from CD8+T-cells, and ADREG5 signals most likely from other cells in the metastatic microenvironment not explored here. CD4+T-cells receive signals from Tregs through the VCAM pathway, as well as signals along the CD45, CCL and ITGB2 pathways. CD8+T-cells are influenced by LCK signals from Tregs, and additional signals along the CD99 and MHC-I signalling pathways. Finally, we compared the overall signalling roles of CD8 and CD4+T-cells in CRCPT and LM. Our analysis revealed that in PT, CD8+T-cells are prominent senders, whereas the CD4+T-cells are prominent receivers. In CRCLM, these roles are strikingly reversed (Fig.4C).

**Fig.4.**
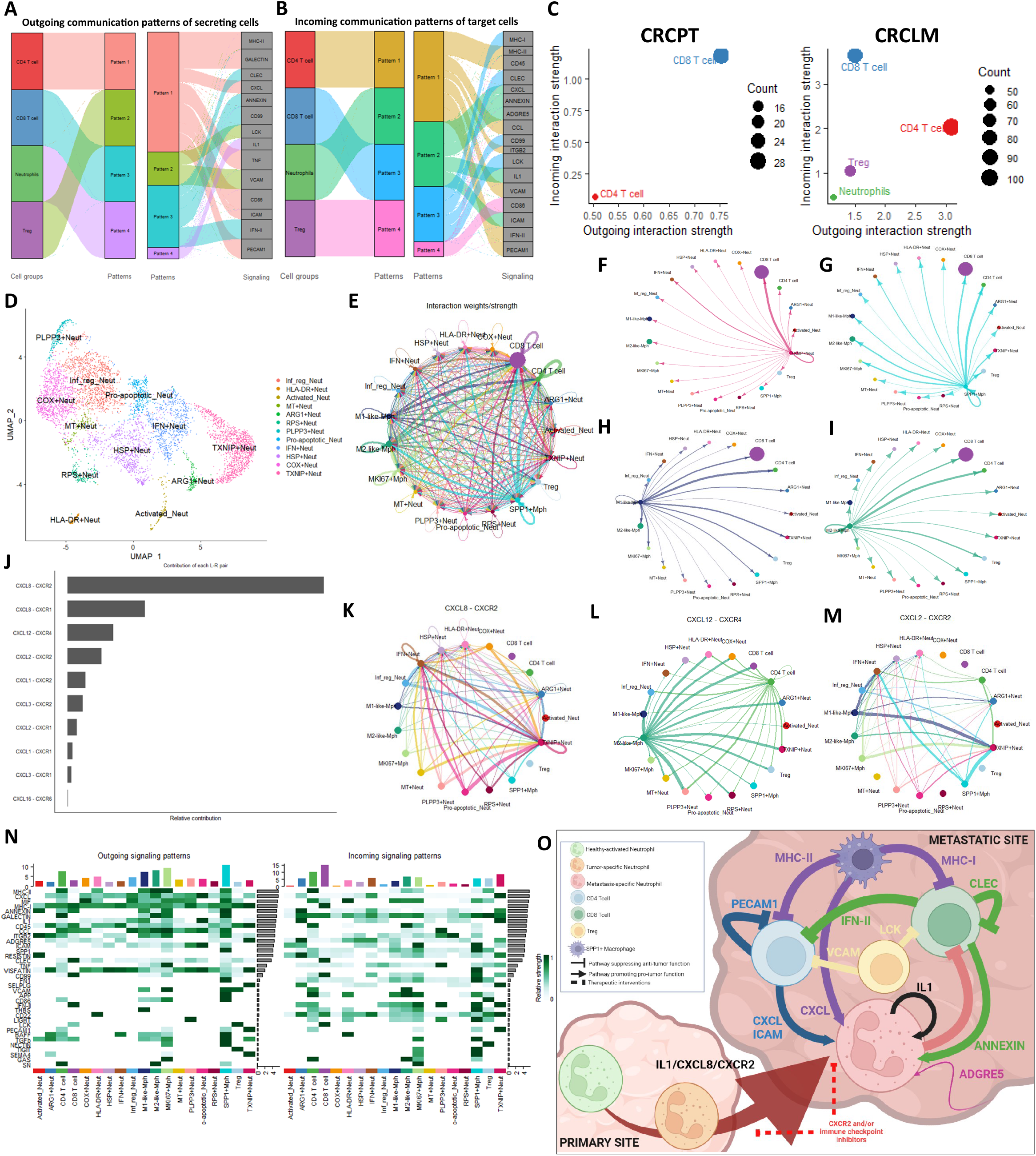
Signalling patterns in CRCLM. (A,B) Outgoing and incoming signalling patterns in CRCLM. © Cellular roles as dominant senders (sources) and receivers (targets) in CRCPT and LM. (D)UMAP plot of neutrophil subtypes in CRCLM. (E) Interaction strength of the global communication patterns between neutrophil, T-cell and macrophage subtypes. (F-I) Outgoing signal strengths from TXNIP+ neutrophils, SPP1+ macrophages, M1- and M2-like macrophages respectively. (J) The contribution of each L-R pair to the overall CXCL signalling pathway amongst macrophage, neutrophil and T-cell subtypes. (K-M) Visualization of the cell-cell communication patterns mediated by the most significant L-R pairs in CXCL pathway. (N) Heatmap showing significant outgoing and incoming signalling patterns in all communication pathways with dominant sender and receiver immune cell subtypes. (O) Summary of findings. IL1/CXCL8/CXCR2 axis drives neutrophil progression from health to cancer and malignancy. Several signalling pathways between neutrophils and other immune cells come to play to foster an immunosuppressive, metastatic niche. Targeting different chemokines in along the IL1/CXCL8/CXCR2 will allow future stratification of treatments into targeting different neutrophil subtypes at early and late stages of cancer. Figure created by biorender.com.

### Macrophages communicate with T-cells and TXNIP+Neutrophils through MHC and CXCL pathways in CRCLM

We then characterised the neutrophil subtypes in CRCLM to investigate their communication patterns with T-cell and macrophage populations from the same dataset, focusing on the immunosuppressive TXNIP+Neutrophils and SPP1+macrophages(16). We identified 11 additional neutrophil subtypes in CRCLM(Fig. 4D, Supplementary Tables S7 and 13): Inflammation regulatory (Inf_reg) neutrophils expressing genes important for inflammatory regulation (*TMG2, CCL4,CCL3 and PI3*), COX+ neutrophils expressing glycolysis genes (*DYNLL1, COX20 and ENO1*), IFN+ neutrophils enriched for interferon response genes (*ISG15, IFIT3, IFIT1 and IFITM3*), ARG1+ neutrophils expressing canonical neutrophil markers (*MMP9 and S100A12*) together with the T-cell suppressive markers *ARG1* and *TXNIP*, HLA-DR+ neutrophils expressing several genes from the human leukocyte antigen (HLA) family (*HLA-DRB1, HLA-DRA*), activated neutrophils enriched for markers of neutrophil activation (*DEFA3, CAMP, LTF, MMP8*) and HSP+, MT+ and RPS+ neutrophils highly expressing heat shock, mitochondrial and ribosomal proteins, respectively and PLPP3+ neutrophils expressing genes that converge on JAK-STAT and EGFR signalling (*PLPP3, FNIP2,PLIN2,SNAPC1, CSTB, CTSD,VEGFA*) and has been reported in other ScRNA-seq studies of neutrophils(13,30).

Analysis of the communication network between neutrophil subtypes, T-cells and macrophages confirmed the recipient role of CD8+T-cells in the CRCLM microenvironment (Fig. 4E) with signals from TXNIP+ neutrophils specifically targeting CD8+ T-cells(Fig.4F). Macrophages exhibited diverse communication patters, interacting with all other immune cell subtypes with stronger interactions with CD4+ and CD8+T-cells. This was observed for all macrophage subtypes (Fig.4G-I). The significant L-R interactions with highest communication probabilities from SPP1+, proliferating MKI67+, M1- and M2-like macrophages targeting CD4+ and CD8+T-cells were through the MHC-II and MHC-I pathways, respectively (Supplementary Fig. S5A-D). Both MKI67+ and SPP1+ macrophages exhibited stronger communication probabilities with M1-and M2-like macrophages along the MIF-(CD74/CXCR4) and MIF-(CD74/CXCR2) axes (Fig.S5C,D). Both M1-like and SPP1+ macrophages showed the highest communication probability with TXNIP+ neutrophils through the CXCL8-CXCR2 L-R interaction(Fig.S5A,D).Further analysis of the CXCL pathway identified CXCL8-CXCR2 as the major L-R interaction (Fig.4J), with additional outgoing signals from other neutrophil subtypes targeting the TXNIP+ population (Fig.4K). M2-like macrophages communicate with all immune cell populations investigated here through CXCL12-CXCR4 interactions (Fig.4L) whereas signals from both M1-like and SPP1+ macrohages target TXNIP+ and IFN+ neutrophils through the CXCL2-CXCR2 interactions (Fig.4M). Finally, analysis of the aggregated cell-cell communication network from all signalling pathways identified both macrophages and CD4+ T-cells as dominant senders in CRCLM (Fig. 4N). Collectively, our data confirm the importance of the CXCL8/CXCR2 axis in immune cell interactions and highlight the dominant roles of macrophages and CD4+ T-cells within the immunosuppressive CRCLM environment.

## DISCUSSION

There is a need to identify novel targets for therapy in advanced CRC to circumvent resistance to current treatments(31). Here we have explored neutrophil and T-cell single cell transcriptomic profiles to assess differences in neutrophil phenotypes in health, primary tumours and metastases and gain insight into how the microenvironment is regulated. The major interactions between the different immune cell populations in CRCLM are summarised in Fig. 4O.

We demonstrate that in health, neutrophils exhibit different transcriptomic signatures according to the tissue they are derived from. This is also true in cancer, where we demonstrated 2 distinct transcriptomic signatures are observed in PT. In metastases, tumour associated neutrophils exhibit the same signatures as the primary site, however, these populations are heterogenous, encompassing additional, distinct transcriptomic changes. Our findings support the role of the tumour microenvironment in recruiting and transforming neutrophils into more immunosuppressive phenotypes (32).

Recent studies have highlighted the roles of opposing neutrophil phenotypes, anti tumorigenic N1 or pro-tumorigenic N2, that exacerbate the progression of cancer depending on their prevalence(11), however, with advances in understanding of plasticity of neutrophils these states appear oversimplified, with neutrophils likely responsive to both the tissue of residence and microenvironmental signalling of the tumour and associated stromal cells. These populations were largely defined based on their function and no cell surface markers have been identified thus far to differentiate between the two(33). In this study, we established the transcriptomic signatures of two distinct neutrophil subtypes in PTs: Healthy enriched and Tumour-enriched neutrophil subtypes, which are conserved across species and across different cancers.

Neutrophils within CRCLM include a subset enriched for genes implicated in T-cell expansion, survival and activation(25,27). This metastatic-specific signature identified in human CRCLM was equally present in a murine CRCLM Bulk-RNAseq dataset generated in our laboratory. Through the TNF pathway, tumour associated neutrophils induce CD8+Tcell apoptosis, further exacerbating their immunosuppressive phenotype(34). Moreover, S100A8 expressing neutrophils facilitate metastasis through the suppression of CD8+T-cells(35). Here, we show an upregulation of TNF signalling in metastatic neutrophils, concomitant with the overexpression of S100A8 and G0S2, implicated in the positive regulation of apoptosis. As such, we propose the presence of a metastasis-specific neutrophil subtype that specifically targets T-cells in order to suppress them.

Our pseudo-time analysis suggests a developmental trajectory of neutrophils that progresses from the healthy subtype to the tumour-specific population and finally a metastasis-specific population; a lineage that is largely driven by IL1B/CXCL8/CXCR2 axis. We show that tumour-associated neutrophils not only respond to IL1B/CXCR2 in their environment but equally signal through IL1B and CXCR2 in an autocrine fashion. The IL1B/CXCL8/CXCR2 axis has been implicated in several tumour types and plays a role in neutrophil recruitment(5,36). The genetic ablation of CXCR2 in mice eliminates tumour accumulation and enhances T-cell infiltration and function(37). Moreover, targeting CXCR2+ immunosuppressive neutrophils, either independently or in combination with additional treatments, enhances anti-tumour immune activity; specifically, that of CD8+T-cells, and reduces tumour burden across different cancer types(38). This supports the presence of a CXCR2+ T-cell supressing neutrophil subset in CRCLM, the elimination of which can enhance T-cell infiltration and function.

It is important to account for the tumour stage when considering the tumour-suppressive effect of targeting CXCR2. We propose that targeting IL1B independently or in combination with CXCR2, could be more favourable at earlier stages, where it may hinder the progression of neutrophils towards the tumour-specific subtype, permitting re-education of neutrophils to a tumour-killing phenotype, in addition to permitting an opportunity for other tumour-directed therapies. Late-stage interventions could target the CXCR2+ T-cell suppressive neutrophil subtypes through utilising CXCR2 antagonists in combination with immune checkpoint inhibitors to counteract T-cell exhaustion. Our findings also implicate the TNF pathway in CRCLM associated neutrophil populations, as well as LTB, complement system and antigen presentation pathways in CD4+T-cells from the same tissue, highlighting the potential of harnessing these aspects of the tumour microenvironment to selectively activate neutrophils for immunotherapy(7).

Finally, we demonstrate that the transcriptomic signature of CD4+T-cells is altered in CRCLM. They are drivers of the signalling network in CRCLM and their interaction with neutrophils, CD8+T-cells and Tregs is essential to mediate immunosuppression as summarised in Fig.4D. CD8+ and CD4+T-cells receive signals along the MHC-I and MHC-II pathways respectively, from macrophages and presumably due to direct interactions with tumour cells. The direct effects of IFN-II on T-cells are largely suppressive(39), thus, we hypothesise that incoming IFN-II signals from CD8+T-cells may drive the suppression of CD4+T-cells in metastasis, specifically the cytotoxic subtype. Amongst the outgoing signalling pathways from CD4+T-cells was GALACTIN. The upregulation of Galectin-9 by IFN-II has an apoptosis-inducing activity in both CD4+ and CD8+T-cells; with CD8+T-cells being more susceptible(40). We identified two autocrine signalling pathways in CD4+ and CD8+T-cells: PECAM1 and CLEC, respectively. The adhesion molecule PECAM1 inhibits T-cell function in mice through the effects of TGF-B(41) and the C-type lectin receptor CLEC-1 negatively regulates antigen cross-presentation by dendritic cells to CD8+T-cells (42), supporting our hypothesis of diminished T-cell activity in the metastatic environment.

We demonstrate that neutrophils receive immunosuppressive signals from both CD4+ and CD8+T-cells. ANNEXIN signalling elicits pro-invasive and pro-tumoral properties in a number of cancers, whereby neutrophil micro-vesicles enriched in Annexin-A1 and TGF-B are immunosuppressive(43). ICAM1 expression immobilises neutrophils and enhances their migration and infiltration(44). Several L-R interactions in the CXCL pathway were between neutrophils, macrophages and CD4+T-cells, suggesting an additional role of CD4+T-cells and macrophages in driving the immunosuppressive neutrophil phenotype. Autocrine IL1B-IL1R2 and CXCL8-CXCL2 interactions within neutrophils support our pseudo-time analysis. Neutrophils are recipients to ADGRE5 signalling, which has a role in tumour invasion and metastasis(45), potentially reflecting tumour-neutrophil interactions. We show that Tregs receive CD86 signals, which upon their engagement with CTLA-4 receptor hamper the antigen presenting ability of antigen presenting cells to activate T-cells(46). They primarily suppress CD4+ and CD8+T-cells via the VCAM and LCK pathways, respectively. VCAM1 is essential for T-cell extravasation and the Src-kinase LCK plays a critical role in initiating and regulating T-cell receptor signalling, whereby LCK inhibition selectively depletes effector Tregs and increases memory CD8+T-cells(47).

In conclusion, there exist two neutrophil transcriptomic subtypes that predominate PTs and are conserved across human and mouse cancers. We propose a developmental trajectory progressing from healthy neutrophils towards a tumour-specific subtype in PTs, with heterogenous expression profiles of neutrophils present within metastases, however, a T-cell suppressive neutrophil lineage can be identified in CRCLM that specifically interacts with CD8+T-cells. This lineage is largely driven by the IL1B/CXCL8/CXCR2 axis. The metastatic niche further fosters an immunosuppressive environment, through the interplay between neutrophils, macrophages, CD8+T-cells, CD4+T-cells and Tregs, with CD4+T-cells and macrophages being dominant signal senders and regulators of the immunosuppressive microenvironment. As such, these interactions, and their timings should be considered when developing future immunotherapy trials in CRCLM.

## Statement of significance

We identify two recurring neutrophil populations and demonstrate their staged evolution from health to malignancy through the IL1B/CXCL8/CXCR2 axis, allowing for immunotherapeutic neutrophil-targeting approaches to counteract immunosuppressive subtypes that emerge in metastasis.

## ACKNOWLEDGEMENTS

We would like to thank Ms. Selena McCafferty from Biorepository Research for facilitating the procurement of human CRCLM tissue for Project #602 when available. The authors would like to acknowledge and thank the McNab family for generous financial support for this project.

## AUTHORS’ CONTRIBUTIONS

RF: Conceptualization, Data generation, Data curation and acquisition, Formal Analysis, Investigation, Software, Validation, Writing – original draft; MW: Data curation and acquisition, Methodology, Visualization, Writing – review & editing; MM: Data curation and acquisition, Methodology; XC-L: Data curation and acquisition; AML: Writing – review & editing, Visualization; JF: Writing – review & editing; KG: Data curation, Methodology, Software; CN: Methodology, Resources; KK: Methodology, Resources; RJ: Writing – review & editing; AD.C: Writing – review & editing; OJ.S: Funding acquisition, Project administration, Supervision; CW.S: Conceptualization, Funding acquisition, Project administration, Supervision, Writing – review & editing.

## DATA AVAILIBILITY

The data generated in this study are available within the article and its supplementary data files. All relevant code has been deposited on Github (https://github.com/ranafetit/NeutrophilCharacterisation). Data analysed from publicly available datasets were obtained from Gene Expression Omnibus (GEO) database and National Omics Encyclopaedia as listed in Table1. Data generated independently from GEM and transplant models of CRC in this study are available from Zenodo at (https://doi.org/10.5281/zenodo.8133995). Any other data generated in this study are available upon request from the corresponding author.

