## Supplementary figures and legends for "Characterising neutrophil subtypes in cancer using human and murine single-cell RNA sequencing datasets"

### Fig. S1

**A**

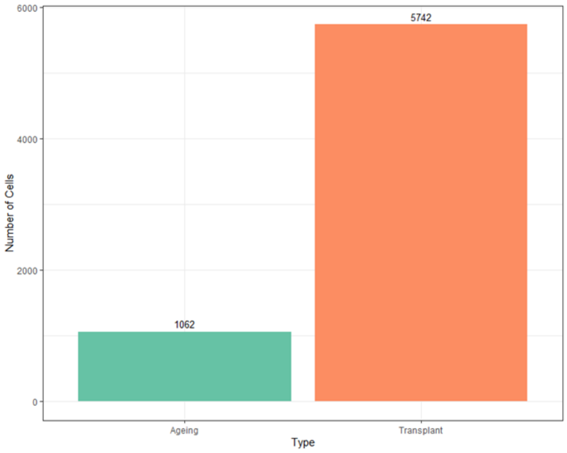

**B**

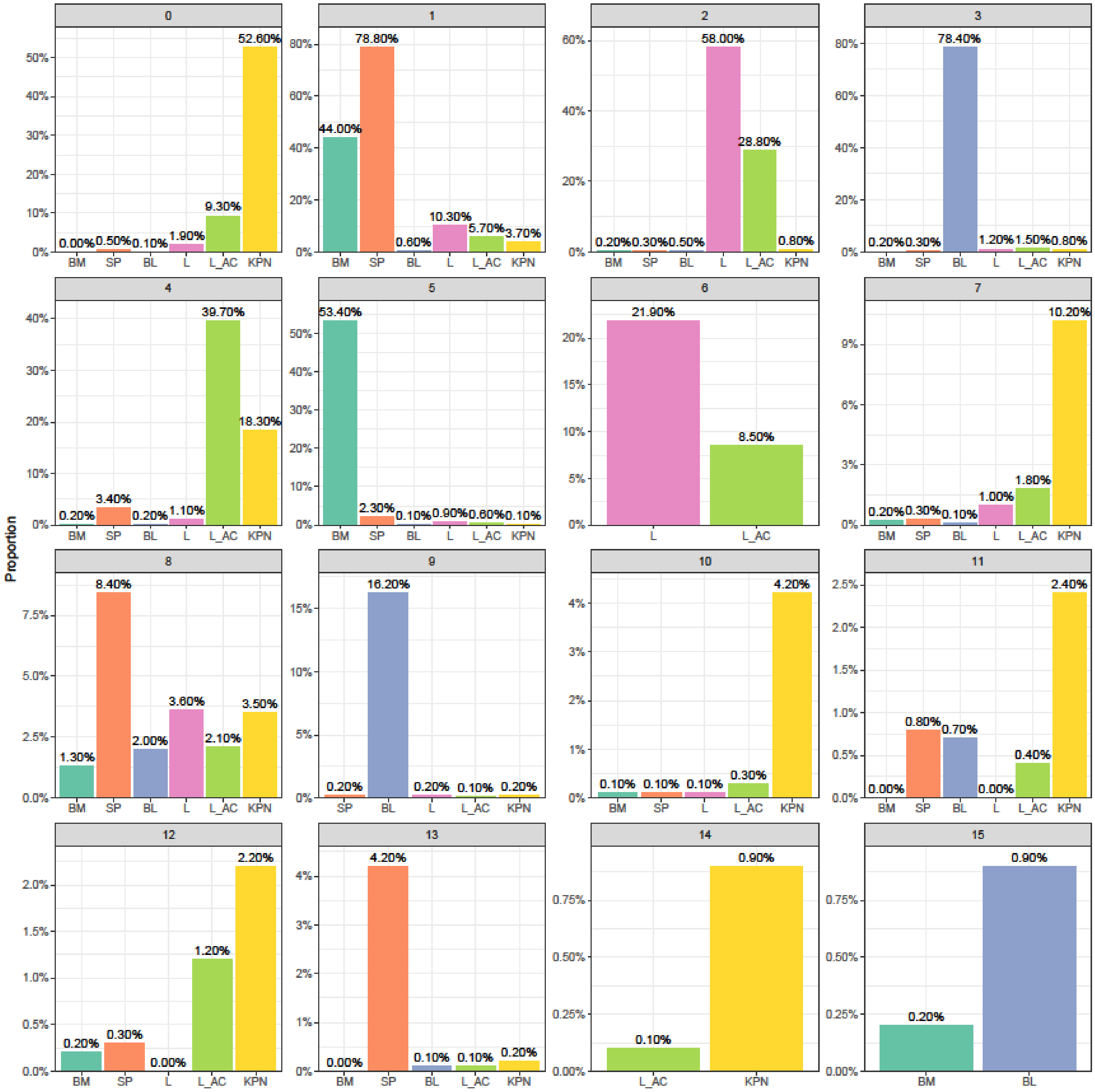

**Fig.S1. Neutrophil proportions in KPN CRC models and in integrated mouse neutrophil dataset.** (A) Proportion of neutrophils derived from aged GEM and transplant models of CRC. (B) Proportion of clusters (from figure 1B) in every tissue type in integrated mouse neutrophil dataset. BM: healthy bone marrow, SP: healthy spleen, BL: healthy blood, L: healthy lung, L_AC: lung adenocarcinoma, KPN: colorectal cancer with *Kras,Trp53* and Notch mutations.

Fig. S2

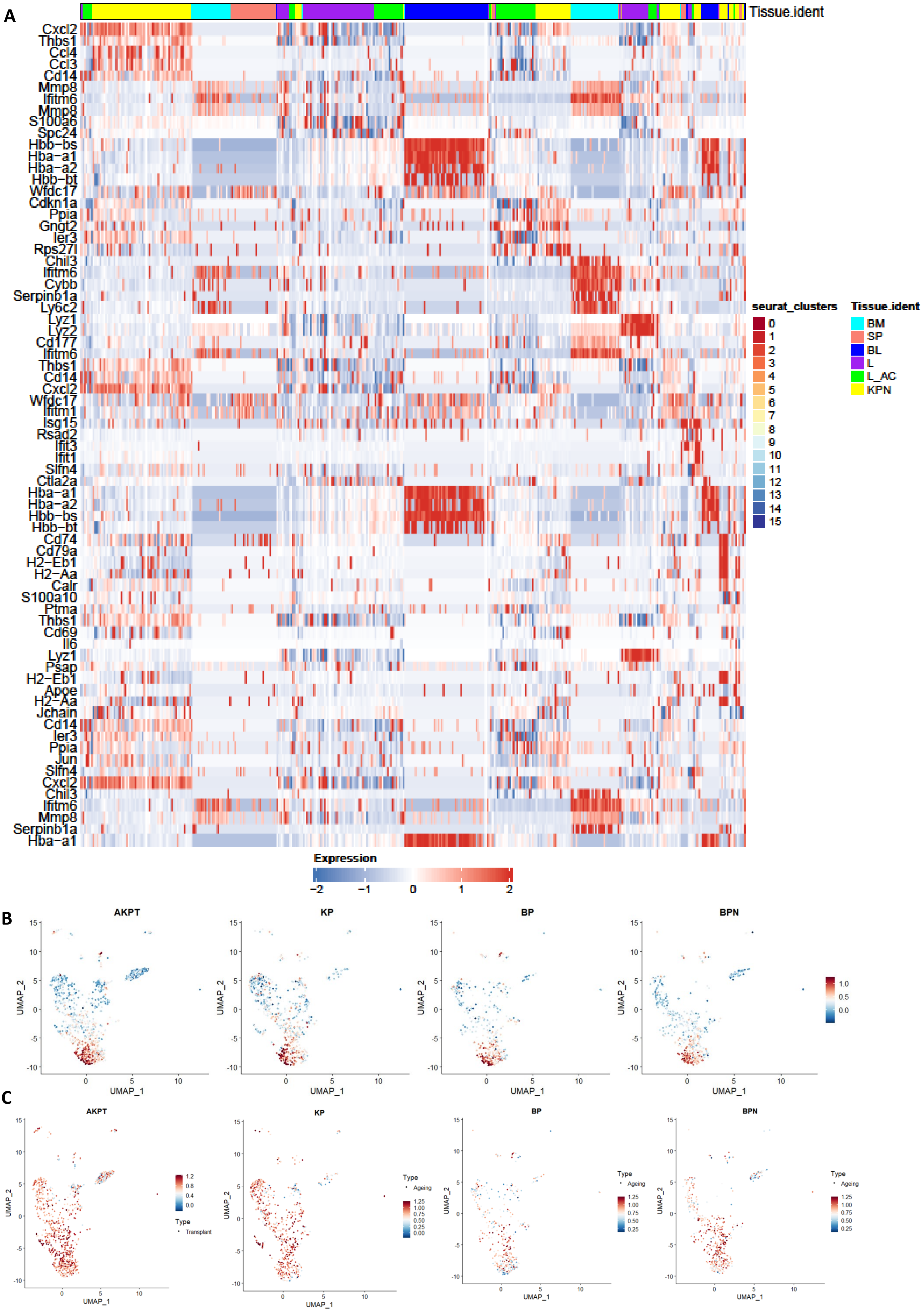
A
seurat_clusters
Tissue.ident
Cxcl2
Thbs1
Ccl4
Ccl3
Cd14
Mmp8
Ifitm6
Mmp8
S100a6
Spc24
Hbb-bs
Hba-a1
Hba-a2
Hbb-bt
Wfdc17
Cdkn1a
Ppia
Gngt2
Ier3
Rps27l
Chil3
Ifitm6
Cybb
Serpinb1a
Ly6c2
Lyz1
Lyz2
Cd177
Ifitm6
Thbs1
Cd14
Cxcl2
Wfdc17
Ifitm1
Isg15
Rsad2
Ifit3
Ifit1
Slfn4
Ctla2a
Hba-a1
Hba-a2
Hbb-bs
Hbb-bt
Cd74
Cd79a
H2-Eb1
H2-Aa
Calr
S100a10
Ptma
Thbs1
Cd69
Il6
Lyz1
Psap
H2-Eb1
Apoe
H2-Aa
Jchain
Cd14
Ier3
Ppia
Jun
Slfn4
Cxcl2
Chil3
Ifitm6
Mmp8
Serpinb1a
Hba-a1
seurat_clusters
Tissue.ident
BM
SP
BL
L
L_AC
KPN
Expression
−2
−1
0
1
2
B
AKPT
KP
BP
BPN
UMAP_1
UMAP_2
1.0
0.5
0.0
C
AKPT
KP
BP
BPN
UMAP_1
UMAP_2
1.2
0.8
0.4
0.0
Type
Transplant
Ageing
1.25
1.00
0.75
0.50
0.25
0.00

**Fig.S2. Gene expression and neutrophil signatures in neutrophils from healthy and tumour tissue.** (A) Heatmap showing the top 5 markers of neutrophil clusters in the integrated mouse dataset. Top bar is coloured according to Seurat cluster, bottom bar is coloured according to tissue identity. (B,C) Healthy and Tumour-specific neutrophil signatures are equally present in both GEM and transplant models of CRC listed in Table2.

**Fig. S3**

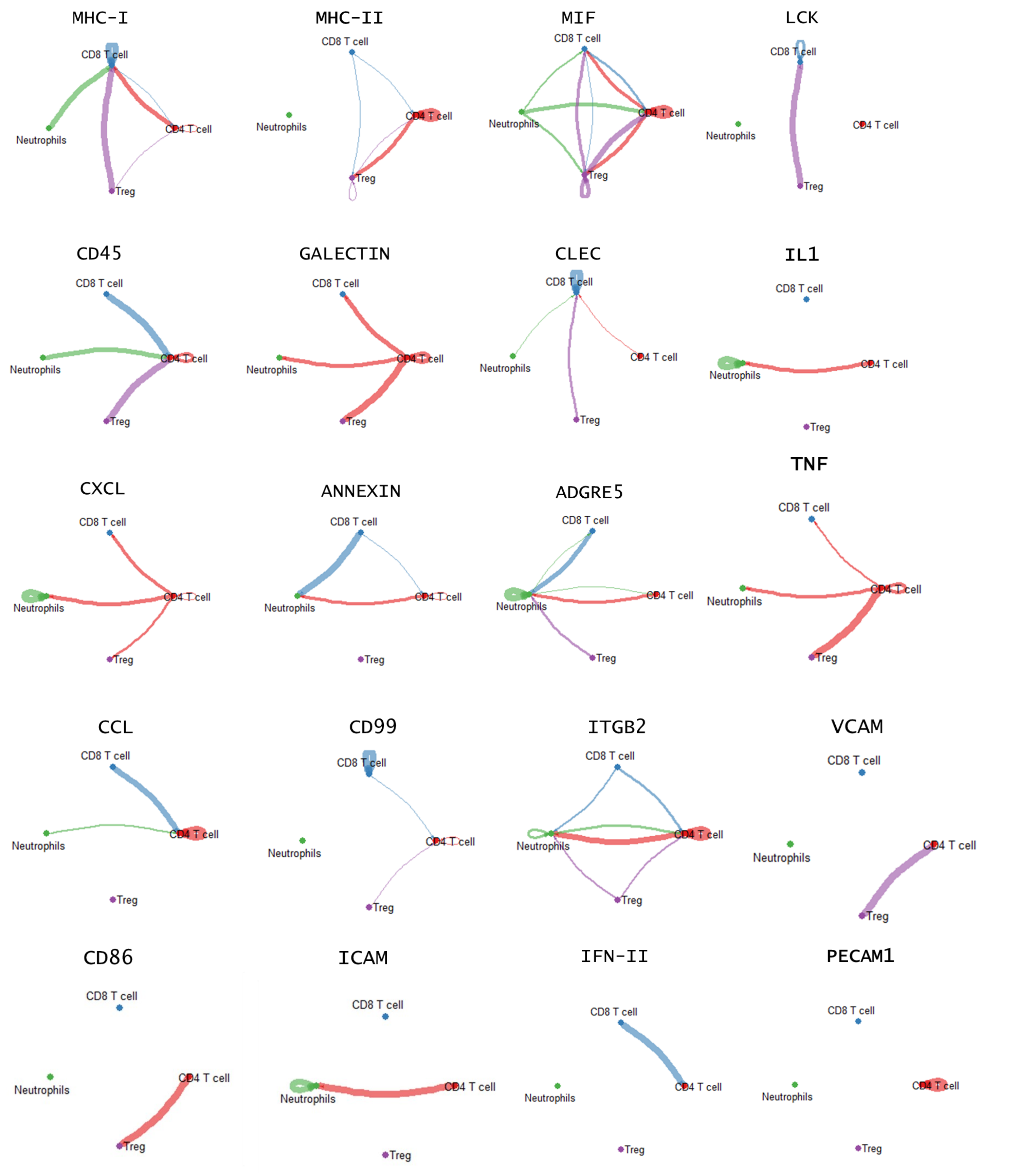

**Fig.S3. cell-cell communication networks at a signalling pathway level.**

Chord plots showing the interaction strengths between the different immune populations in CRCLM across the 20 signalling pathways with significant communications.

**Fig. S4**

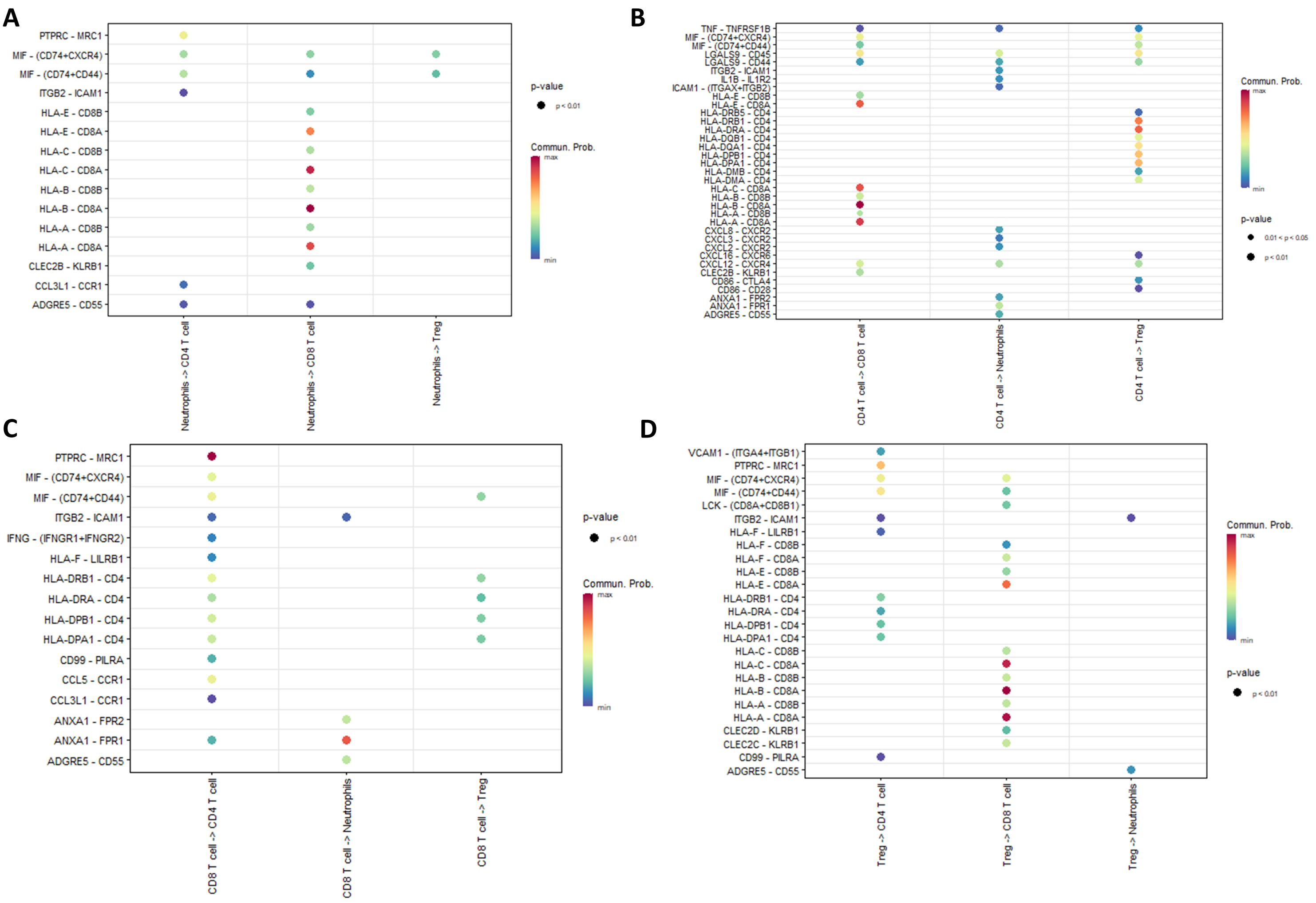

**Fig.S4. L-R interactions from the significant signalling pathways that mediate neutrophil and T-cell communication in CRCLM.**

(A-D) Bubble plots showing significant L-R interactions and their communication probability across the 20 significant pathways from neutrophils, CD4+T-cells, CD8+T-cells and Treg cells to other cell groups.

**Fig. S5**

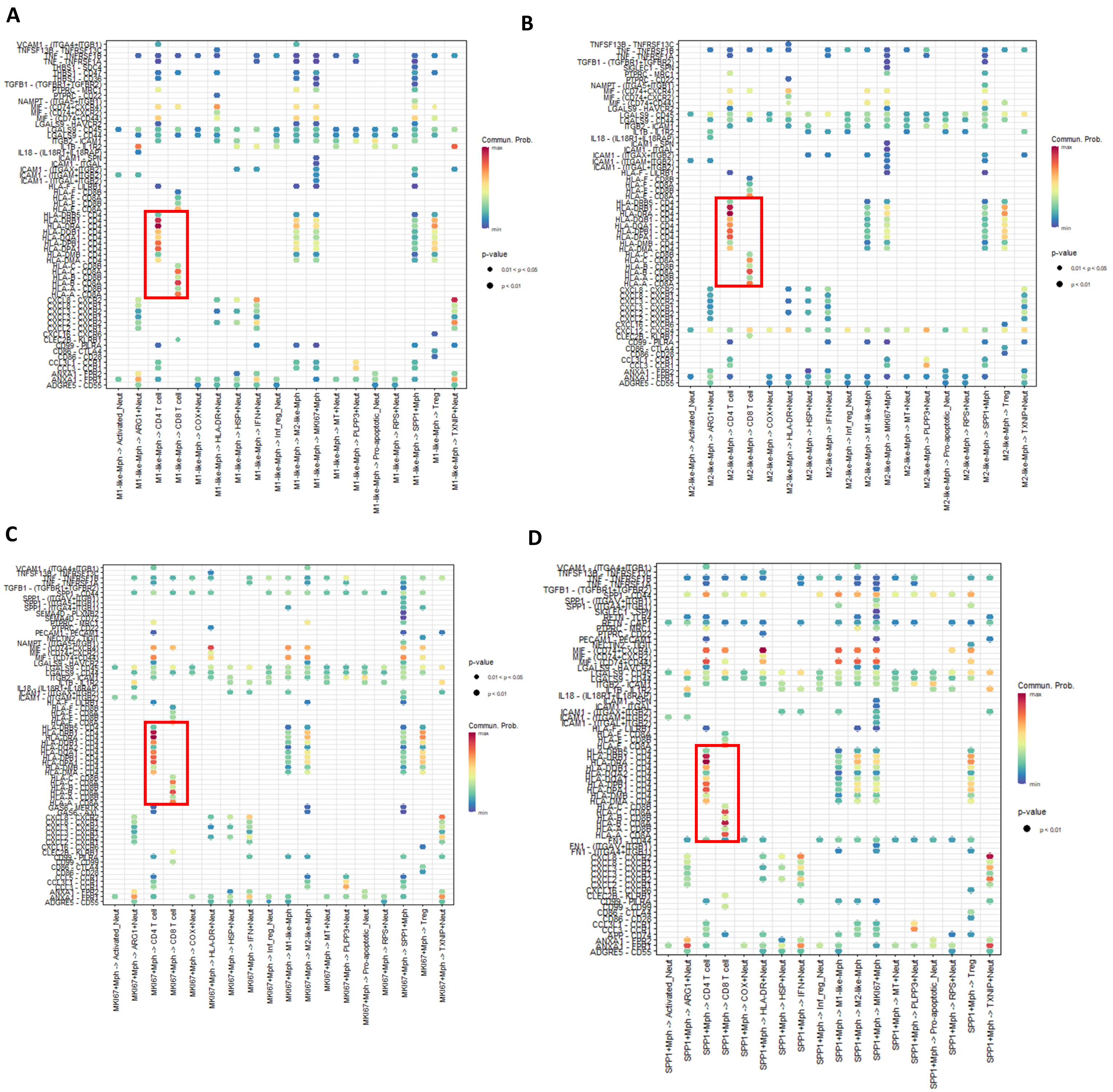

**Fig.S5. L-R interactions from the significant signalling pathways that mediate communication between macrophages and other immune-cell subtypes in CRCLM.**

(A-D) Bubble plots showing significant L-R interactions and their communication probability across the significant pathways from M1-like, M2-like, MKI67+ and SPP1+ macrophages towards other immune-cell subtypes. Red boxes highlight the significant L-R interactions with highest communication probabilities targeting CD4+ and CD8+T-cells through MHC-II and MHC-I pathways.
